# Analysis of the computational strategy of a detailed laminar cortical microcircuit model for solving the image-change-detection task

**DOI:** 10.1101/2021.11.17.469025

**Authors:** Franz Scherr, Wolfgang Maass

## Abstract

The neocortex can be viewed as a tapestry consisting of variations of rather stereotypical local cortical microcircuits. Hence understanding how these microcircuits compute holds the key to understanding brain function. Intense research efforts over several decades have culminated in a detailed model of a generic cortical microcircuit in the primary visual cortex from the Allen Institute. We are presenting here methods and first results for understanding computational properties of this largescale data-based model. We show that it can solve a standard image-change-detection task almost as well as the living brain. Furthermore, we unravel the computational strategy of the model and elucidate the computational role of diverse subtypes of neurons. Altogether this work demonstrates the feasibility and scientific potential of a methodology based on close interaction of detailed data and large-scale computer modelling for understanding brain function.

## 1 Introduction

A major insight into brain function was the discovery that the mammalian neocortex is in first approximation a continuous 2D sheet consisting of rather stereotypical cortical microcircuits (Mountcastle 1998; Douglas and Martin 2004; Harris and Shepherd 2015). This architecture offers hope that one can understand brain function by understanding the computational organization of its local processors: cortical microcircuits. The structure of these cortical microcircuits, which are sometimes referred to as cortical columns, appears to be highly preserved from mouse to human. Different types of neurons are arranged on roughly 6 parallel sheets or laminae, forming synaptic connections primarily to nearby neurons within the same or other laminae. Hence both its spatial organization and its units, consisting of a fairly large set of neuron types with diverse response properties, mark salient differences to generic recurrent neural network models that are commonly considered in computational neuroscience, and abstracted into artificial neural networks in modern AI.

The large number of genetically, morphologically, and electrophysiologically different neuron types in the mammalian neocortex, as well as the technical difficulty to probe the efficacy of synaptic connections between every pair of neuron types, have made it difficult to determine the generic structure of cortical microcircuits. But intense research during the past three decades (Mountcastle 1998; Thomson and Lamy 2007; Markram et al. 2015) has recently culminated in a detailed cortical microcircuit model (Billeh et al. 2020) for area V1 in mouse, see Figure 1, to which we will simply refer as the Billeh model. Now the challenge arises to relate the structure of this model to its computational function. One obstacle is that many parameter values, such as the strength of synaptic connections between individual neurons, are still missing, and are not likely to be determined in the near future through experimental work. We have quite a number of experimental data about the average strengths of synaptic connections between the main neuron types, especially if their somata have little distance. Quite a bit of this general statistical knowledge has entered the heuristic setting of weights in the Billeh model. But the higher order moments of these weight distributions remain unknown. This knowledge gap makes it hard to relate the structure of microcircuit models to their computational function, since the latter is likely to arise largely from the composition and alignment of individual synaptic weights. This situation is comparable with that in artificial neural networks, where even perfect knowledge of the histogram of synaptic weights in a trained network provides almost no insight into what it has been trained for. The alignment of synaptic weights arises in the brain through a host of plasticity processes, which create correlations and higher order dependencies that are likely to define the computational role of individual neurons and local network motifs within the larger network. Hence, in order to investigate computational capabilities of the cortical microcircuit model of Billeh et al. (2020) one needs to examine the results of aligning or optimizing individual synaptic weights for concrete network computations. One commonly refers to this alignment process as training of the network model.

**Figure 1:**
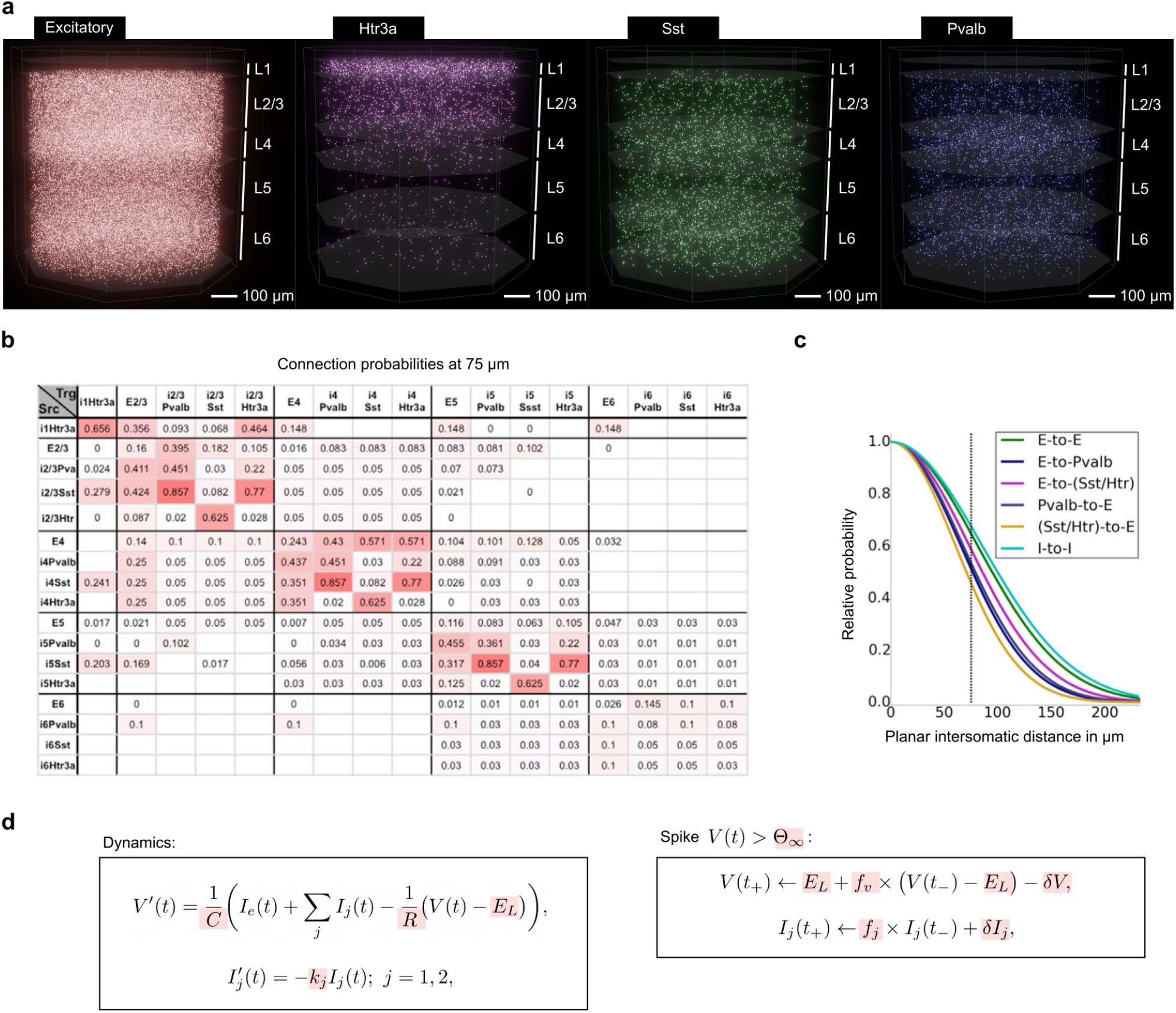
Overview of the data-based cortical microcircuit model of Billeh et al. (2020). **a)** Visualization of the locations of the 51,978 neurons in the model from Billeh et al. (2020), split into excitatory and 3 major classes of inhibitory neurons (Htr3a, Ssst, Pvalb). Only the 5 laminar sheets L1, L2/3, L4, L5, L6 are distinguished in this model. **b)** Base connection probabilities between 17 different types of neurons that result if one takes laminar locations of neurons from the previously shown 4 major classes into account. **c)** Scaling of connection probabilities in dependence of somatic distance for different types of connections. The probability of a synaptic connection results by multiplying the base connection probability for the two neuron types involved with this scaling function. **d)** Main equations and parameters that govern the dynamics of internal variables and spiking activity of the employed GLIF_3_ neuron model. Highlighted parameters stem from experimental data as described in Billeh et al. (2020), giving rise to 111 different neuron models in the microcircuit according to experimental data from the cell database of the Allen Brain Atlas.

We report here methods and first results of this research strategy. We have adapted the synaptic weights of the Billeh model through stochastic gradient descent to support a particular type of network computation: the image-change-detection task. This task has frequently been used in biological experiments on mice (Garrett et al. 2020; Joshua H. Siegle et al. 2021): The subject receives a long sequence of natural images, with short time gaps in between where just a uniformly gray screen is shown. Training the Billeh model to solve this computational task is for several reasons not straightforward:

i. If one models the neurons as leaky integrate-and-fire (LIF) neurons, this neuron model is not differentiable.
ii. The point neuron models of Billeh et al. (2020) pose additional challenges: They are generalized LIF neurons, more precisely, GLIF_3_ models that contain two additional internal variables which model slower dynamics processes such as after-spike currents, as found in biological neurons (Teeter et al. 2018).
iii. Even the core model of Billeh et al. (2020) that we consider is fairly large: It consists of 51,978 neurons. But gradients need to be computed very fast nevertheless, since successful stochastic gradient descent training typically requires the computation of gradients for the whole network for very large numbers of input presentations (trials).

We found that recently proposed approximations of stochastic gradient descent for recurrent networks of LIF neurons (Bellec et al. 2018) can be efficiently adapted to work also for GLIF_3_ neuron models. Furthermore, we show that very efficient software (TensorFlow (Martin Abadi et al. 2015)) and computer hardware (GPUs), that have been designed to support fast training of deep neural networks in machine learning, can be adapted to train large and biologically detailed models for deep neural networks of the brain, such as the model of Billeh et al. (2020).

We demonstrate the potential of this research strategy by training the Billeh model for the image-change-detection task, and then “opening up the black box” (Sussillo and Barak 2013) of the trained model in order to elucidate how it carries out this quite demanding network computation. We show that by applying reverse engineering methods to the computer model that can at present not be applied to the living brain one can understand how the diverse neuron types of the model can collaborate to carry out this network computation.

## 2 Results

### 2.1 A data-based laminar microcircuit model can solve the image-change-detection task

The microcircuit model of Billeh et al. (2020) provides a major advance, since it is based on an extensive body of experiments at the Allen Institute that were all directed at one brain area, V1, in one species, mouse, see Figure 1a, b and c. More precisely, we have used the “core” part of the point-neuron version of their model, since simulations of the detailed biophysical version require too much compute time. But the point neuron version provides already a major advance over previous models because it is based on 17 different data-based neuron types (listed in each row and column of Figure 1b). These are further split into 111 different variations based on response profiles of individual neurons from the Allen Brain Atlas (Allen Institute 2018) to which GLIF_3_ neurons have been fitted.

We trained the Billeh model to solve the image-change detection task through backpropagation-through-time (BPTT). We expanded the BPTT method of Bellec et al. (2018) for LIF neurons so that it could be applied to GLIF_3_ neuron models. We did not allow synaptic weights to change their sign, thereby preserving Dale’s law. We included a regularization term similarly as in Bellec et al. (2020) in the loss function for gradient descent in order to keep the firing activity of the network in a biologically realistic sparse firing regime. As a result, the distribution of firing rates stayed close to the biological data from Joshua H. Siegle et al. (2021) and to the rate distribution before training, see Supplementary Figure S2. In particular, the average firing rate after training was 3.86 Hz. Hence the model computed in an energy-efficient sparse firing regime. The distribution of synaptic weights changed during training only little for synaptic connections between excitatory neurons, and weights generally became stronger for synaptic connections from and to inhibitory neurons, see Supplementary Figure S3.

The Billeh model received for the image-change-detection task, like the subjects of the biological experiments (Garrett et al. 2020; Joshua H. Siegle et al. 2021), a sequence of natural images, interleaved by short phases where a gray screen was presented as visual input, see Figure 2a, b. These natural images were first processed by the model for the LGN (lateral geniculate nucleus) of Billeh et al. (2020), producing input currents to neurons of the microcircuit model in a retinotopic and lamina-specific manner as in Billeh et al. (2020), see Figure 2c. The task of the subjects was to report whenever the most recently presented image differed from the previous one (Figure 2a). The model was trained to report within a response window of 50 ms length that started 50 ms after image offset through increased firing of a population of excitatory neurons in L5 if the image was different from the preceding one. Since we do not have experimental data about the identity of the readout neurons that extract the network decision and project it to other brain areas, we randomly selected in our model 60 excitatory neurons from a sphere with a diameter of 170 microns within layer 5 (see Figure 2e) to produce together a number of spikes that transcended a decision threshold (see bottom row of Figure 3a) whenever the image had changed. This modelling assumption appears to be reasonable since pyramidal cells on layer 5 are typically viewed as readout neurons from a laminar cortical microcircuit, reporting network decision in particular to subcortical structures (Harris and Shepherd 2015).

**Figure 2:**
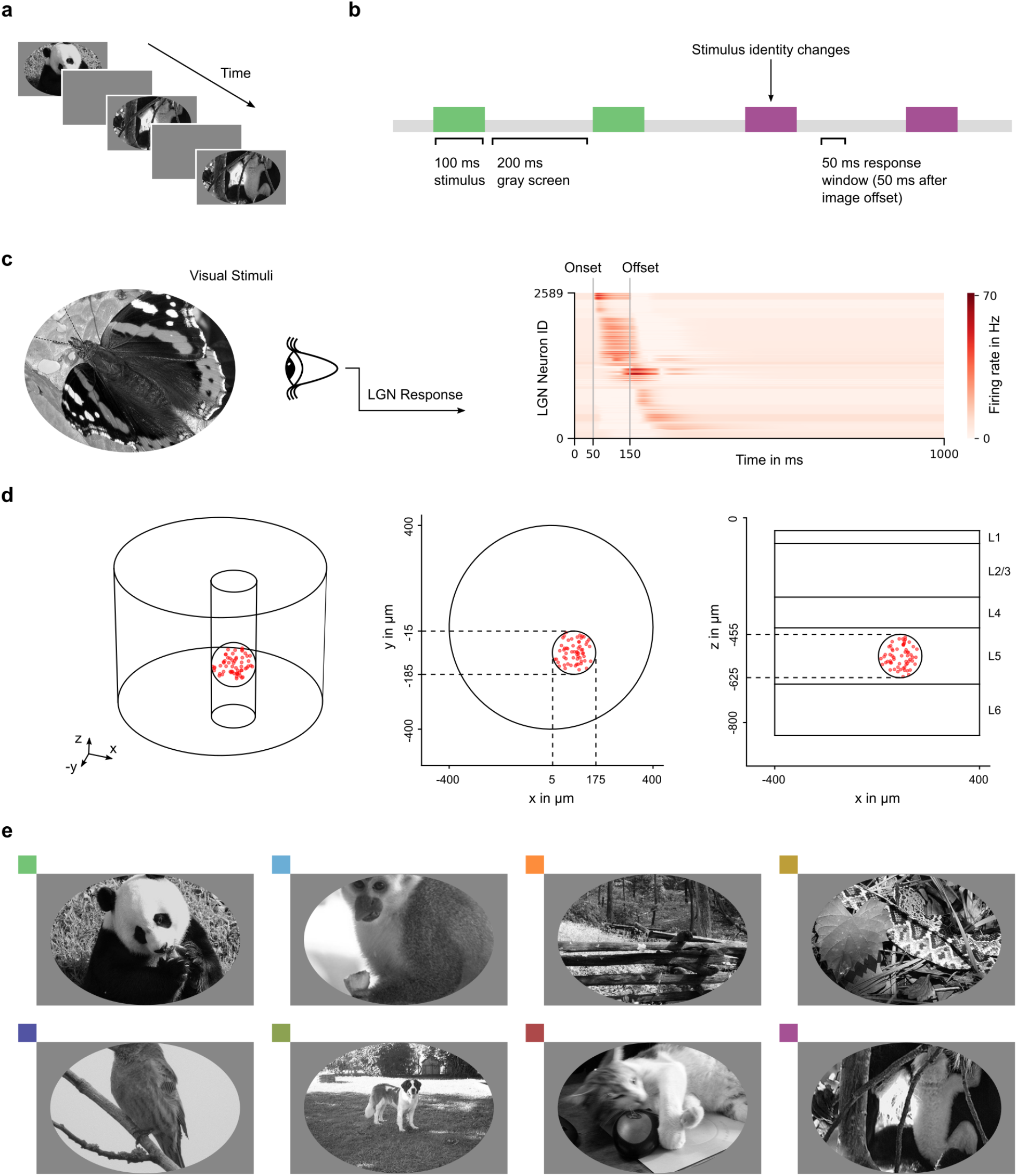
Image-change-detection task. **a)** Schematic sequence of visual stimuli in the task. Images sequences are presented to the model, interleaved by delays of gray screen **b)** Diagram explaining the temporal structure of the task. The model has to report if a different image than the previous one is shown during the 50 ms response window. **c)** A model of LGN encodes the visual stimuli into a temporal response of LGN neurons that are associated to a particular location in visual field (The responses of the LGN neurons shown here were sorted by the timing of peak activation). This response serves as input to the data-based model of V1, and is injected as currents instead of sampled Poisson spike trains. **d)** A random selection of 60 excitatory neurons in layer 5 (within a sphere of 170 microns in diameter) constitutes the readout. These neurons report a positive decision collectively by high firing rates. **e)** For testing, images for the image-change-detection task were drawn from this set of 8 images that are not used during training.

**Figure 3:**
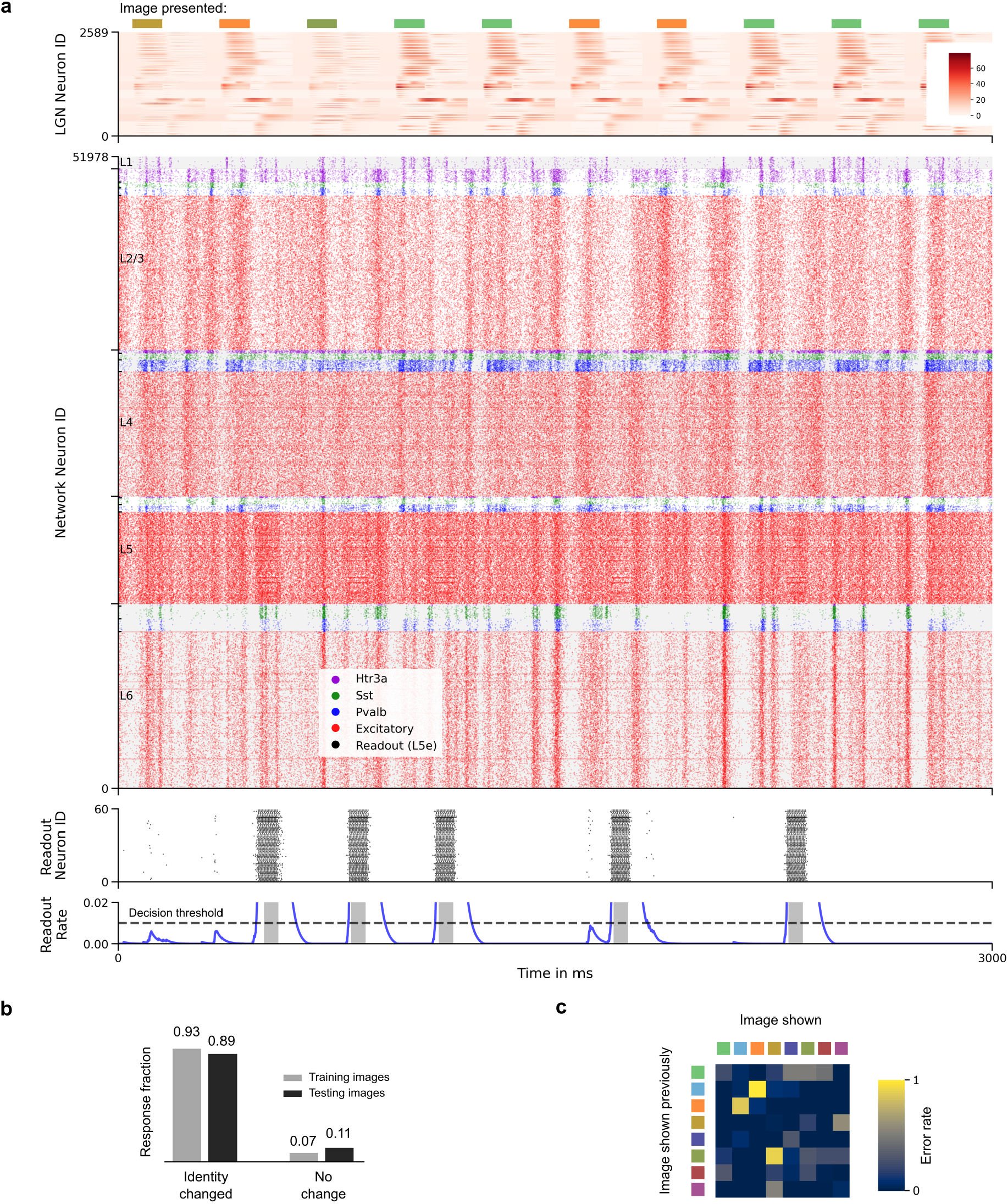
Sample of network activity during task performance after training. **a)** Visual stimuli composed of a sequence of images, interleaved by delays of gray screen, are converted to a response pattern using the model of LGN (top row). This input is injected in a current-based manner using the data-based LGN to V1 connections into the neurons of the model, resulting in network activity (middle row, neuron types and layers are separated but the order within these groups is randomized). Whenever the presented image is different to the previous one shown, the 60 readout neurons in layer 5 (L5) report it by high firing rates during a response window of 50 ms (penultimate row). The rate of this readout can be estimated using an exponential sliding window (blue curves). Its temporal evolution is shown along with the decision threshold (last row). **b)** After the training procedure, the model could perform the task for images used during training, but also for new, unseen images. For those testing images (see Figure 2e) the model was able to report the changed identity in 89% of all cases, while it wrongly reported a change in 11% of cases where the image identity did not change. **c)** Transition-specific errors. For certain types of transitions between different images the model incurs more errors than otherwise, especially for the images marked light blue and orange.

We randomly selected a pool of 48 natural images from the Imagenet dataset (Deng et al. 2009) that we used as network inputs. We used 40 of them for training, similar to the biological experiments of Garrett et al. (2020). Task performance was evaluated both for the 40 images used for training, and for the other 8 images. The model achieved after training a high performance for this task, see Figure 3b, that lies in the same range as the performance achieved by mice (Garrett et al. 2020). Importantly, the trained model was able to generalize very well, achieving -like the subjects of Garrett et al. (2020)-almost the same performance for images that were not used during training. Hence the model has a general computational competence that is not constrained to particular images. In the subsequent sections we will “open the black box” and unravel the strategy that the laminar microcircuit model uses for the processing of these new images.

We also analyzed in which cases errors arise most often, see Figure 3c. It can be inferred that the model mostly has problems due to a confusion of the images associated with the colors orange and light blue, regardless of which came first. Correspondingly, the distance-preserving low-dimensional projection of network states in Figure 4c shows that the network states are the least separated when these two images have been processed.

**Figure 4:**
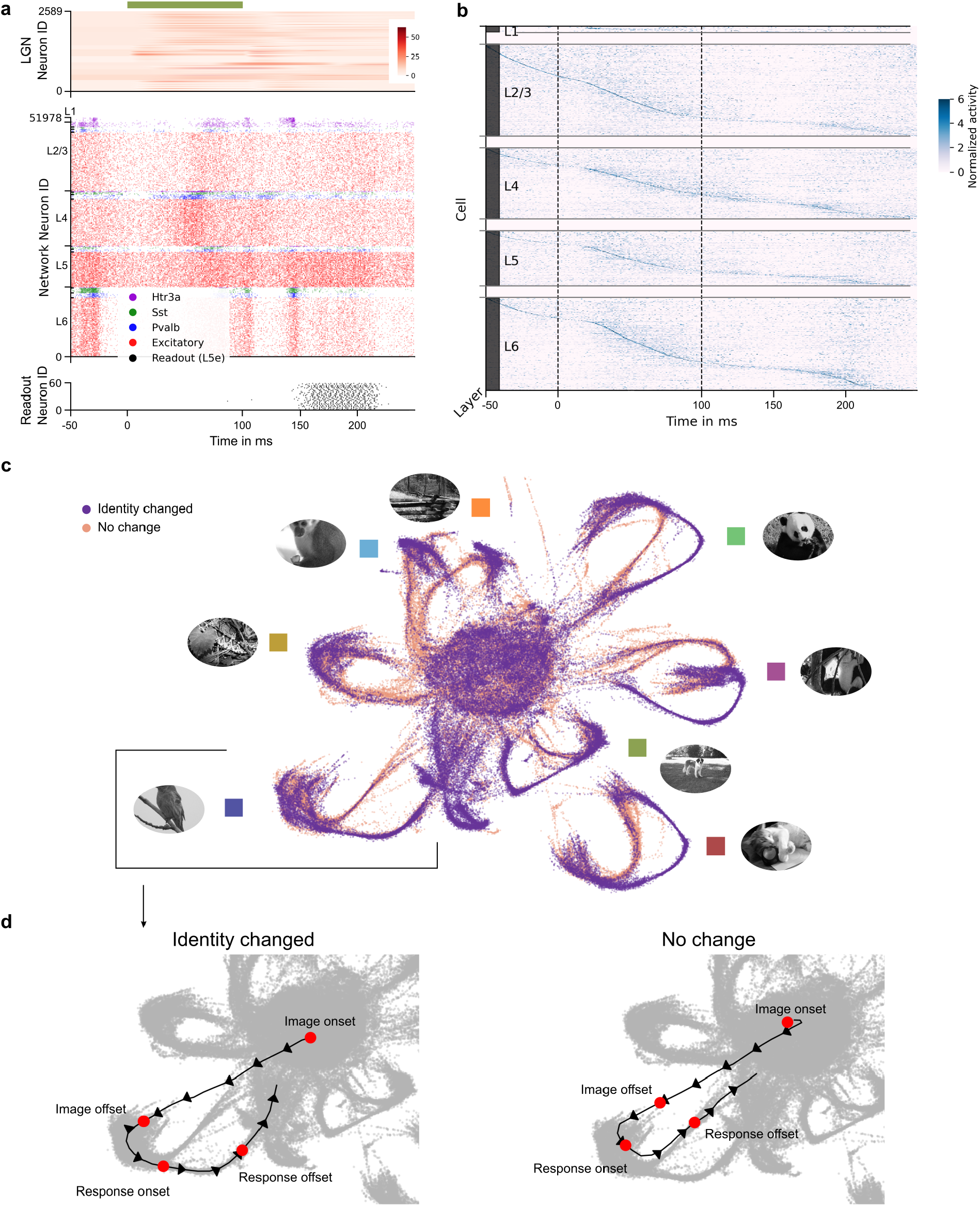
Global perspective of network computations as dynamical system. **a)** Spike raster of the network in response to an image presentation, on a finer time scale than Figure 3a. **b)** Preferred time of activity of each neuron is shown. Different laminae can be seen to compute in parallel, rather than sequentially. The timing of neural activity is less stereotypical during the response window, since we average here over change and no-change conditions. **c)** Low dimensional embedding of network activity using UMAP (McInnes, Healy, and Melville 2018). Each dot represents the network activity at a particular ms during the processing of an image and the generation of the network decision. The embedding was obtained by considering spike trains of all neurons during 150 seconds of task performance. These spike trains were subjected to an exponential filtering procedure using a kernel with a time constant of 20 ms. The filtered spike trains were subsequently projected to their 50 most salient principal components (PCA), which were then embedded into 2D space by an application of UMAP **d)** Averaged network embedding trajectories for a single image, both for the case where the preceding image was different and where it was the same. These trajectories emerge by averaging the low-dimensional projections of network trajectories.

### 2.2 Linking network dynamics and network computation

A direct link between network dynamics and network computation -i.e., behavior of the organism-was exhibited for c-elegans through Ca-imaging data by Kato et al. (2015). They found that most neurons participate in the examined behavior, in spite of their numerous differences in spatial collection and genetically encoded neuron type. Furthermore, they demonstrated that different behaviors, which correspond in our model to computations with different network decisions (change or no-change), can be clearly decoded from low-dimensional projections of the temporal evolution of the high-dimensional vector formed by the states of individual neurons. We wondered whether similar links can be drawn for network computations in our model, in spite of numerous deviations from the paradigm of Kato et al. (2015):

i. computer model of a brain network versus in-vivo recordings
ii. mammalian neocortex versus the nervous system of c-elegans
iii. visual perception task versus motor behavior
iv. state vectors defined by of spiking activity of neurons versus derivatives of the internal Ca-dynamics of neurons
v. 51,978 dimensional state vectors versus maximally 131-dimensional recorded state vectors of celegans.

We now focus on the network dynamics during processing of an image and during the subsequent response window, as shown in Figure 4a. We first address the question whether most neurons participate in such a generic computation, and if so, whether there is a rule when they usually become active. Since our model exhibits a fairly large variety of network responses to the same image, due to additional random inputs summarized as “rest of the brain” (Billeh et al. 2020), we considered in Figure 4b trial averaged activity: We normalized the activity of each neuron over the time course under consideration, and plotted its averaged activity. This normalized activity allows us to plot the time at which each neuron tends to become most active, independently of their overall activity level. We sorted all neurons according to the time of the peak of their activity relative to image onset, see Figure 4b. This analysis suggests that most neurons participate in the network computation, each at a preferred time. In particular we can discern two classes of neurons: Those that prefer to become active during an image presentation, and those that become more active after an image presentation.

We further performed an embedding of the network activity during task performance using the 2D projection UMAP (McInnes, Healy, and Melville 2018) of the 51,978-dimensional network states that result from the spiking activity of its 51,978 neurons, see Figure 4c. More precisely, we applied an exponential filter with a time constant of 20 ms to the spike output of each neuron for 8 new images that had not been used during training. We then discarded all but the 50 most important principal components of these network states, which were then embedded into 2D space by UMAP. This analysis reveals in Figure 4c that the network undergoes during image processing a directional low-dimensional dynamics, which can be seen as the backbone of the network computation into which the computational processing of each neuron is embedded. Furthermore, each stimulus (image) and network decision (behavior) produces a bundle of trajectories in the network dynamics that stay in general well-separated, except for the case of the two images marked blue and orange in Figure 2e, for which a change between them is less reliably detected by the network according to Figure 3c. Hence one sees here a direct link between the structure of the network dynamics and its computational performance.

A refined temporal evolution of network states is shown in Figure 4d for cases where always the same image was presented, but the preceding image was either different or the same. The trajectories of network states are almost the same during the presentation time of the image, no matter whether it had occurred already just before or not. But a clear bifurcation of network states is visible at the onset of the response window.

Altogether one sees that the dynamics and computational organization of the nervous system of c-elegans exhibits numerous parallels with our trained data-based model for area V1 in mouse.

### 2.3 Where and when does the network decision emerge?

The low-dimensional projection of global network states in Figure 4 showed a bifurcation of network trajectories for the change/no-change condition after the offset of an image. But where and when does information about the decision emerge in the network? To answer this question we analyze in Figure 5a the temporal and spatial organization of information about the network decision. The first information about the network decision arises in the firing activity of neurons during the time window from 50 to 100 ms after the onset of an image. This time interval is of particular interest because information from a new image reaches the microcircuit model about 50 ms after image onset. Hence this time interval is the earliest possible one where any neuron could possibly disseminate information through its spikes regarding the question whether the current image is the same as the preceding one. Figure 5a shows that there exist in fact neurons, primarily in L4 and L5, whose spikes contain already during this earliest possible phase information about the subsequent network decision, that is created 100 ms later, during the response window that lasts from 150 to 200 ms after image onset. Figure 5a indicates the locations of those neurons that have during this earliest phase the largest MI with the subsequent network decision. Figure 5b shows the spiking activity of 7 sample neurons whose firing rates during this time interval has substantial MI with the network decision. They were among those 12 that had the highest MI, but we do not show those 7 with the largest MI in order to allow inclusion of examples for Sst and Pvalb neurons. Most of these neurons are excitatory, but also Sst and Pvalb neurons (color code for neuron types as in Figure 3a) are included. Whereas most of these neurons had a higher firing rate for the change condition, one of the two examples for Pvalb neurons (blue spikes) among the selected 7 neurons uses a lower firing rate in the change condition, the other one a higher firing rate. Also all excitatory neurons among them have during [50, 100] ms a higher firing rate in the change condition.

**Figure 5:**
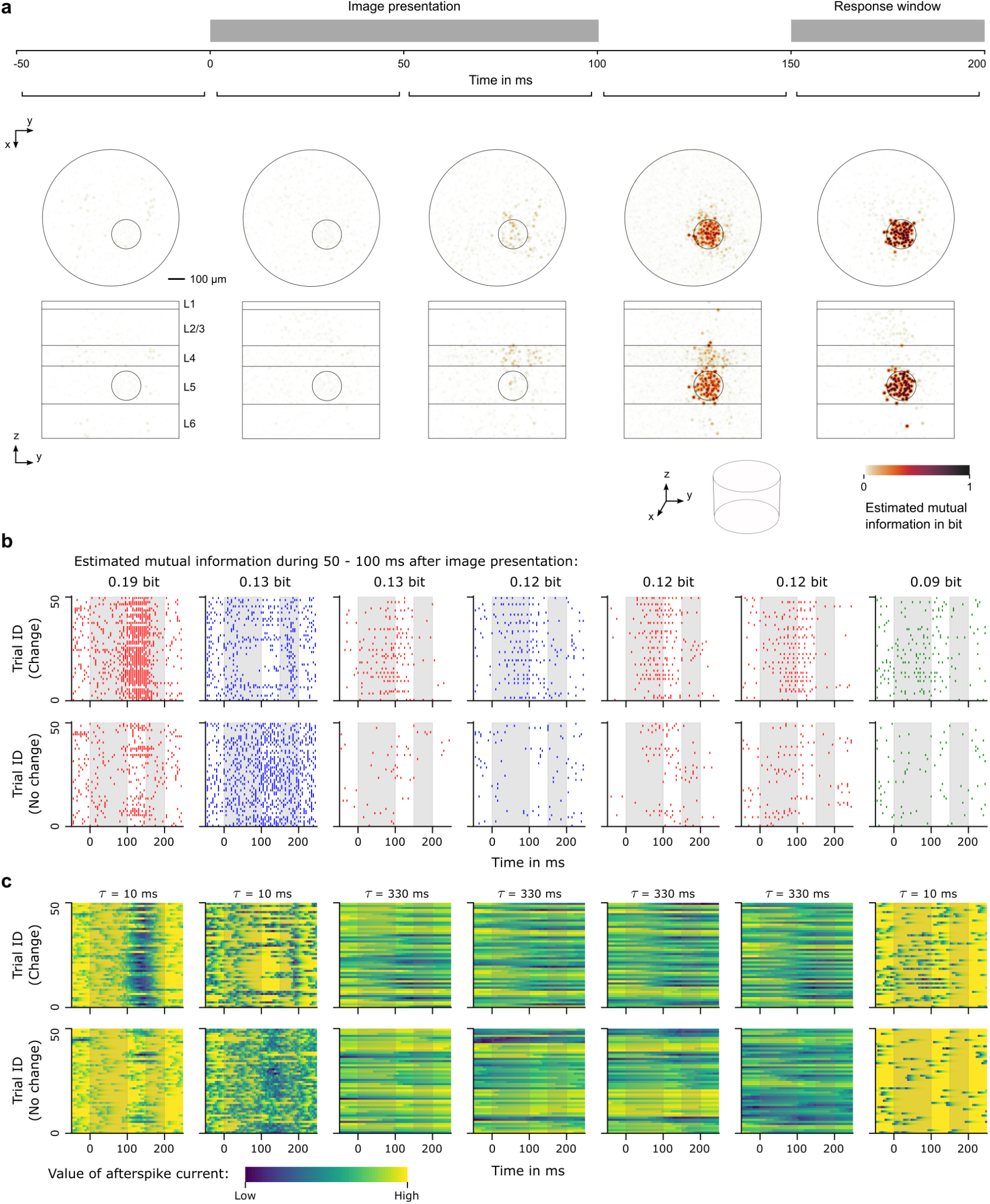
Mutual information between neuron activity and network decision. **a)** The mutual information between activity of single neurons and the change/no-change decision of the network during the response window can be empirically estimated. This was achieved by considering the spike counts of each neuron within 50 ms windows and establishing an empirical joint distribution for spike count and network decision. This was then used to compute the mutual information. For neurons that overlap in the projection from 3D to 2D the maximum value of them is visualized, thereby avoiding that dark points can arise through accumulation of small contributions from several neurons. **b)** Spike trains of neurons with high mutual information during the period from 50 ms to 100 ms after image onset. Trials are separated depending on the change/no-change condition. One sees condition-dependent differences in their firing responses from about 50 ms after image onset. **c)** Visualization of the slowest after-spike current of the same neurons and trials (time constant shown on top).

Figure 5c shows the time course of the slowest internal variable of the underlying GLIF_3_ models of these 7 neurons. The time constants of their slowest internal variable -for an after-spike current-is indicated at the top of each column in Figure 5c. Hence those with large time constants could potentially convey information about the preceding image, whose offset was 200 ms before the onset of the current image. This will be further analyzed in the next subsection.

### 2.4 Neuronal mechanism that triggers the network decision

Figure 5c suggests that long-lasting after-spike currents play an important role in the network computation for image-change-detection since many of those neurons that transmit the earliest information about the subsequent network decision have after-spike currents with large time constants. Figure 6a shows the locations and types of those 20 neurons with a time constant larger than 300 ms whose firing during [50, 100] ms that have the highest MI with the subsequent network decision. All 4 major neuron types occur among them, and they are mostly located in L4, especially at the border to L2/3. In Figure 6b we analyze for four neurons that have large MI the dependence of the value of their slow after-spike current at the onset of the current image, in dependence of the identity of the preceding image. The first column provides an overview of this dependence, showing that the largest current amplitude was specific to a particular identity of the preceding image. The histogram of values of this internal variable at the onset of the present image is shown in the second column. The last column shows that its extreme values (more than 2 std from the mean) are assumed almost exclusively for a particular preceding image. In other words, they use the value of their after-spike currents as working memory for the identity of the preceding image. Hence, if this value of their internal variable can be made observable, i.e., transformed into spiking activity that affects the readout neurons in L5, this working memory could be used to produce a correct network decision. Figure 6c shows that they do in fact play a pivotal role in the production of the network decision: Silencing of these neurons one after the other, starting with those that have the highest MI during [50, 100] ms, degrades the accuracy of the network quite strongly, reaching chance level when all 20 neurons shown in the first panel are silenced. Hence these 20 neurons are causally related to the network decision.

**Figure 6:**
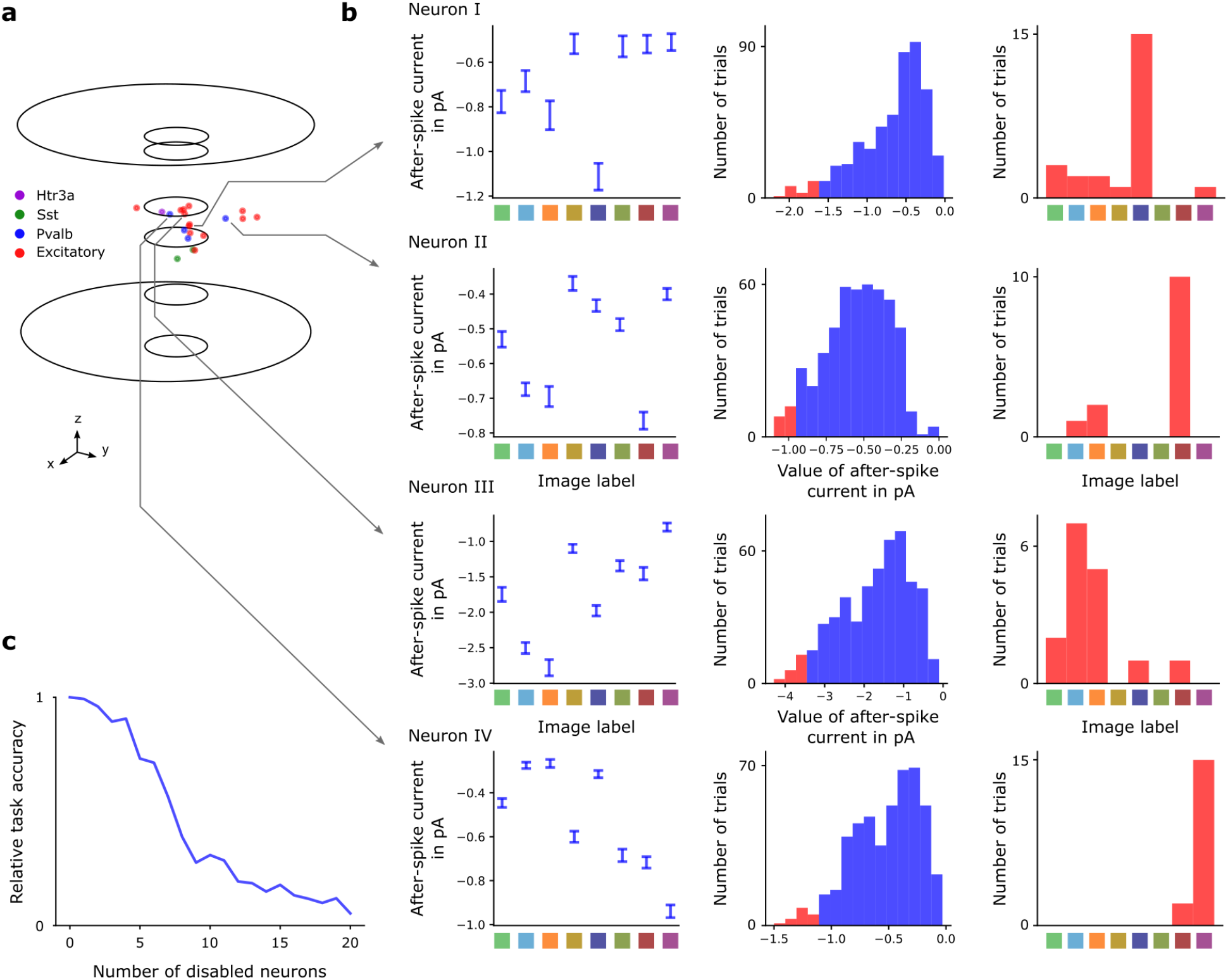
Identification of neurons that trigger the network decisions. **a)** Spatial location and type of those 20 neurons with a large time constant (*>* 300 ms) for after-spike currents that have during [50, 100] ms the largest MI with the network decision (during the response window). **b)** Statistical analysis of the values of their after-spike currents at the onset of a new image, in dependence of the identity of the preceding image (left column). The mean is shown with error bars denoting the std. The analysis of trials where this variable assumed its most negative values (at least 2 std from the mean, colored red) reveals that these neurons are highly selective to the identity of the PRECEDING image: however, since the network had been trained with different images, they appear to select each some generic image feature. This analysis is shown for 4 neurons that were randomly selected among the 20 neurons from panel a) (bottom row). **c)** Verification that the spiking activity of the 20 neurons from panel a) is causal for the network decision. Task performance visibly drops as these neurons are deactivated one after another, from highest to lowest MI of their spiking activity during [50, 100] ms with the network decision.

Finally, we would like to emphasize that the computational analysis in Fig. 5 and 6 was carried out for a new set of images that had not been shown during training of the network. Hence our reverse engineering has unraveled the generic computational mechanism of the network, rather than one that was induced for concrete images.

## 3 Discussion

We have shown that one can train a large and biologically detailed model for a patch of neocortex to carry out a demanding computational task. We have focused here on a task that has also frequently been considered in biological experiments: to report an image change in a sequence of natural images that is interleaved with gray screens. In fact, after training the large-scale model for a patch of area V1 of Billeh et al. (2020) the model reaches for the image-change-detection task the same performance level as the subjects (Garrett et al. 2020). Furthermore, the model is able to solve this task for new sets of images. In other words, it has learned to apply a network algorithm that is generally applicable. Our reverse engineering of the resulting network computation shows that a particular feature of the neuron models in Billeh et al. (2020), that are based on the detailed cell type data of the Allen Brain Atlas, plays a key role in the resulting network computation: The presence of internal variables of neurons that change on a much slower time scale than the membrane potential. These variables, which reflect for example after-spike currents, are usually not considered in computational models for neural networks of the brain. Our hypothesis is that they play an essential role in computations of generic cortical microcircuits, especially for computations where time and delays play an essential role.

We have also shown in Figure 4c and d that a powerful method for the conceptualization and visualization of network computations that have been developed for c-elegans by Kato et al. (2015) can be adapted for cortical microcircuits models to elucidate also in their much higher dimensional space of network states the relation between network dynamics and network function (behavior). Like in their data for c-elegans we see that computational progress in the cortical microcircuit model produces a directional low-dimensional trajectory of network states to which most neurons in the network contribute, in spite of their different types, subtypes, and laminar locations. Trial-to-trial variability gives rise to bundles of such trajectories that need to be well-separated for different network inputs and conditions in order to avoid erroneous network decisions. This dynamical system perspective enables us to understand the global reference framework into which computational contributions of individual neurons, such as those exhibited in Figure 6, are embedded. It also allows us to relate the network dynamics to deficiencies of its computational performance (failure to detect changes between the images marked blue and orange). Altogether, if recordings from mouse V1 support the predictions of our model, this will show that salient aspects of the organization of computations have been preserved from c-elegans to mouse V1, in spite of the obvious differences that we have listed in subsection 2.2.

This work demonstrates the feasibility of a powerful methodology for understanding brain computations: An interplay of detailed biological data and computer simulations of large-scale models that carry out the same computational task as the brain areas from which one records, thereby enabling the generation of detailed hypotheses about the computational organization and underlying neural mechanisms. On the practical level we have shown that software tools (TensorFlow) and computer chips (GPUs) that have been developed to accelerate deep learning applications in AI make this method accessible for many researchers. An obvious next step in this direction is the investigation of distributed computations in several cortical microcircuits. Numerous experimental data suggest that microcircuits of the neocortex carry out computations interactively with microcircuits in other neocortical and subcortical areas, but we do not know much about the organization of these distributed brain computations. In particular, it has been conjectured that working memory function is distributed over several cortical areas, and that working memory for longer time spans is contributed by higher cortical areas. Pyramidal cells in L2/3 of neocortical microcircuits are conjectured to serve as hub for integrating information streams from lower and higher areas. Specifically, we conjecture that an expansion of our model through data-based interconnections with microcircuit models for higher cortical areas will make it possible to solve the image-change-detection task also for longer intermittent periods between image presentations, as in the experiments of Garrett et al. (2020) and Joshua H. Siegle et al. (2021). On the other hand, our results suggest that V1 can solve this task without contributions from higher brain areas for the case of temporal distances of up to 200 ms between successive images. Furthermore they suggest that L2/3 is less essential for this simpler version of the task. In addition, our Figures 5 and 6 point to an important role of L4 for this version of the task. The underlying connectivity data of Billeh et al. (2020) suggest that L2/3 is in fact in a key position for solving this task, because pyramidal cells in L4 have direct synaptic connections to pyramids in L5 with about 75% of the connection probability to L2/3 pyramids. Hence L2/3 is likely to become computationally less relevant in a model where it does not also receive top-down inputs. Furthermore, the data of Billeh et al. (2020) show that many pyramidal cells in L2/L3 and L4 have after-spike currents with long time constants, a feature that is essential for solving the image-change detection task according to our results. This finding is of particular interest in view of previous paradigms for modelling computations in neural networks of the neocortex, which rarely addressed the functional role of neuron diversity and longer time constants of neurons. In contrast, we have demonstrated the feasibility of a more integrated research approach where detailed physiological and anatomical data are directly combined with the analysis of computations in large-scale network models. One nice feature of this approach is that it generates a substantial number of hypotheses that can be experimentally tested since they suggest recordings from particular types of neurons in particular locations of laminar cortical microcircuits.

## 4 Methods

### 4.1 Details of the training procedure

In order to train the model, we considered the following loss function:

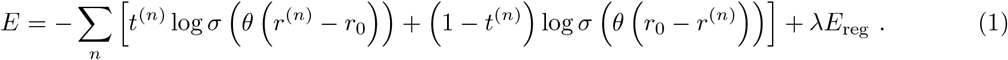

Here, the sum over *n* is organized into chunks of 50 ms and *r*^(*n*)^ denotes the population firing rate of the readout neurons in that time interval. Similarly, *t*^(*n*)^ denotes the target output in that time window, being 1 if a change in image identity should be reported and otherwise 0. The value *r*_0_ = 0.01 denotes a baseline firing rate. The term *λE*_reg_ is a regularization term that penalizes unrealistic membrane voltages as well as unrealistic firing rates. We applied BPTT, backpropagating errors within consecutive time windows of 700 ms length (see Figure 7), and minimized the loss function with respect to the weights between the neurons in the model, and the parameter *θ >* 0. Specifically, we employed 64 GPUs of the JUWELS Booster to carry out this optimization program, where gradients were computed on 128 sequences in parallel. See also Supplementary Figure S1 for an overview of the scaling behavior in the distributed training setup.

**Figure 7:**
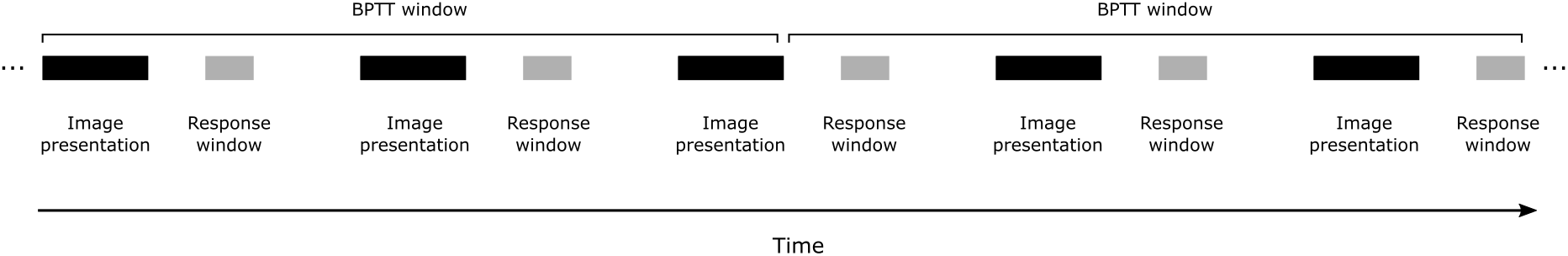
Visualization of BPTT windows during task performance. BPTT is applied to windows of 700 ms, regardless of the alignment to the image presentation.

### 4.2 Software and hardware details

The BPTT training algorithm was coded in TensorFlow, which runs very efficiently on GPUs. A simuation of the Billeh model for 700 ms of biological time and computation of BPTT-gradient through this computation took about 5 s on a fast GPU (NVIDIA A100). This computation had to be iterated 16,000 times in order to achieve high computational performance for the chosen task, which took 23 h of wall clock time on 64 GPUs (see Figure S1 for the speedup resulting from this parallelization).

### 4.3 After-spike currents provide working memory similar to threshold adaptation

It was shown in Bellec et al. (2018) that an adapting threshold enables working memory along the lines of LSTM networks by considering slow internal processes of neurons. In particular, the proposed model was denoted LSNN, and includes neurons that emit a spike *z*(*t*) = *H*(*v*(*t*) − *A*(*t*)) whenever the membrane voltage *v*(*t*) crosses an adaptive threshold *A*(*t*) from below (*H* denotes the Heaviside function). In their case, the adaptive threshold can be written in terms of the filtered spike train of the same neuron

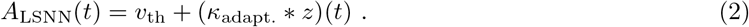

Here, *v*_th_ denotes the baseline threshold, * denotes the convolution operation, and *κ*_adapt._ is a causal, exponential kernel: 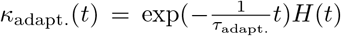, where *τ*_adapt._ is the time constant of adaptation. Hence, the threshold would increase with every emitted spike, and fade back to its baseline thereafter.

The point neuron model of (Billeh et al. 2020) consists of GLIF_3_ neurons (Teeter et al. 2018). These do not include an adaptive threshold but so-called after-spikes currents, which inject current into the membrane after spike emission. The injected current decays according to a specific time constant thereafter. It is argued that if this injected current is negative, it will essentially have a similar effect as an adapting threshold and hence potentially providing also a similar capability for working memory. In fact, one can rewrite the dynamics of GLIF_3_ neurons with an after-spike current in terms of an adapting threshold. In doing so, one can write the dynamics of GLIF_3_ neurons as LSNN neurons but with a different adaptive threshold 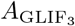 that emerges from an application of 2 filters on the spike train:

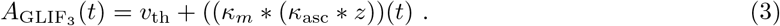

Here, *κ*_*m*_ is a causal, exponential kernel with a time constant of the membrane voltage of the neuron, and *κ*_asc_ is defined using the time constant of the after-spike current. Importantly, this suggests that both models should possess the same capabilities for working memory provided that the time constants of the slower internal processes (adapting thresholds or after-spike currents) are comparable. It also suggests that the working memory that is implemented by the after-spike currents in the GLIF_3_ model reacts slower due to an additional filter.

**Derivation** We will describe the neuron models in terms of differential equations, closely relating to the definition of GLIF in Teeter et al. (2018). Let *R, C* and *I*_*e*_(*t*) denote the membrane resistance, membrane capacitance and input current to the considered neuron respectively. Further, assume that *E*_*L*_ denotes the resting potential of a neuron and *v*_th_ is its baseline threshold.

Using these definitions, one can define the dynamics of the membrane voltage *v*(*t*) and of the adapting threshold *A*_LSNN_(*t*) of a neuron as in Bellec et al. (2018) using the equations:

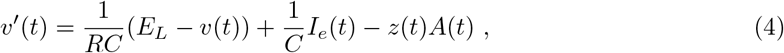

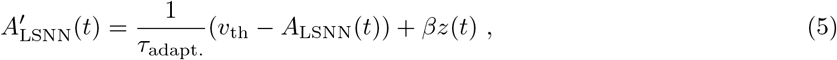

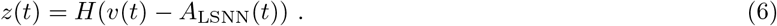

Note that *z*(*t*) is the spike train of the neuron, *H* is the Heaviside function, *τ*_adapt._ is the time constant of threshold adaptation and the parameter *β* scales the impact of threshold adaptation.

In contrast, the GLIF_3_ neuron model, as introduced by Teeter et al. (2018), does not include an adapting threshold but a number of after-spike currents. Consider the case when there is just a single after-spike current *I*_asc_(*t*), then the dynamics of a GLIF_3_ neuron can be expressed by the following equations:

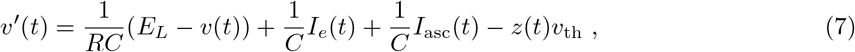

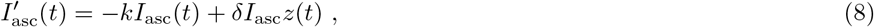

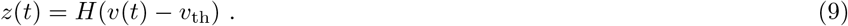

Here, *κ* corresponds to the inverse time constant of the decay of the after-spike current and *δI*_asc_ denotes the increase in after-spike current right after a spike.

It is possible to bring equations (7)-(9) into the form of equations (4)-(6), where we introduce an adapting threshold 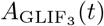 for GLIF_3_ neurons. This allows us to compare and interpret the slower internal mechanisms (threshold adaptation and after-spike currents) on a common ground. For this purpose, we substitute 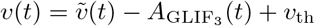 in equations (7)-(9). Equation (7) then becomes:

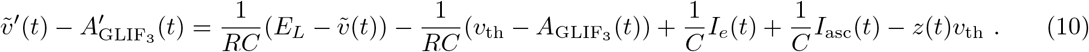

This differential equation can be split in two (which when subtracted from another yield the original one), and thus yield the dynamics of the GLIF_3_ model in terms of an adapting threshold, facilitating a comparison between LSNNs and GLIF_3_:

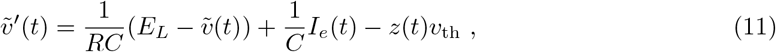

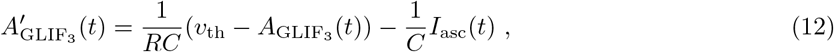

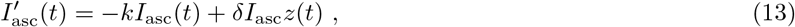

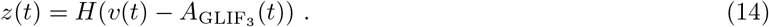

Note that the solution of *A*_LSNN_ is given by:

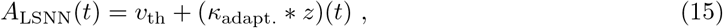

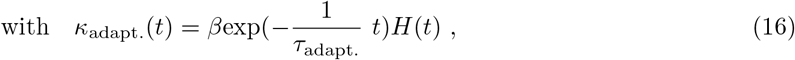

where * denotes the convolution operation. The solution of 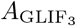, on the other hand, includes an intermediate integration due to *I*_asc_, hence resulting in:

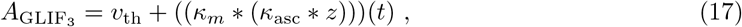

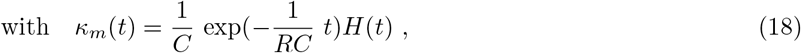

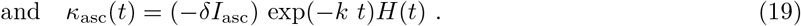

## Acknowledgements

We would like to thank Jason Maclean and Sacha van Albada for helpful discussions, as well as Sandra Diaz for advice and help regarding large-scale computations. This research was partially supported by the Human Brain Project (Grant Agreement number 785907) of the European Union and a grant from Intel. Computations were carried out on the Human Brain Project PCP Pilot Systems at the Jülich Supercomputing Centre, which received co-funding from the European Union (Grant Agreement number 604102).

