## Supplementary figures for "Analysis of the computational strategy of a detailed laminar cortical microcircuit model for solving the image-change-detection task"

November 2021

### 1 Supplementary Figures

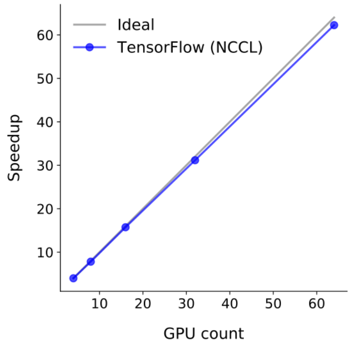


Figure S1: **Scaling properties of the distributed training setup.**

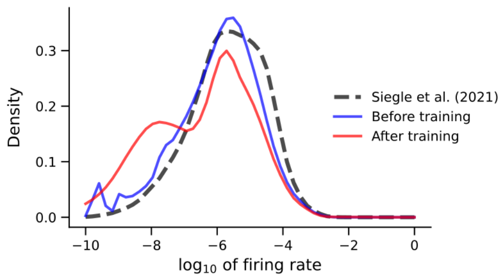


Figure S2: **Firing rate distribution of the Billeh model before and after training, compared with experimental data from (Joshua H. Siegle et al. 2021)**

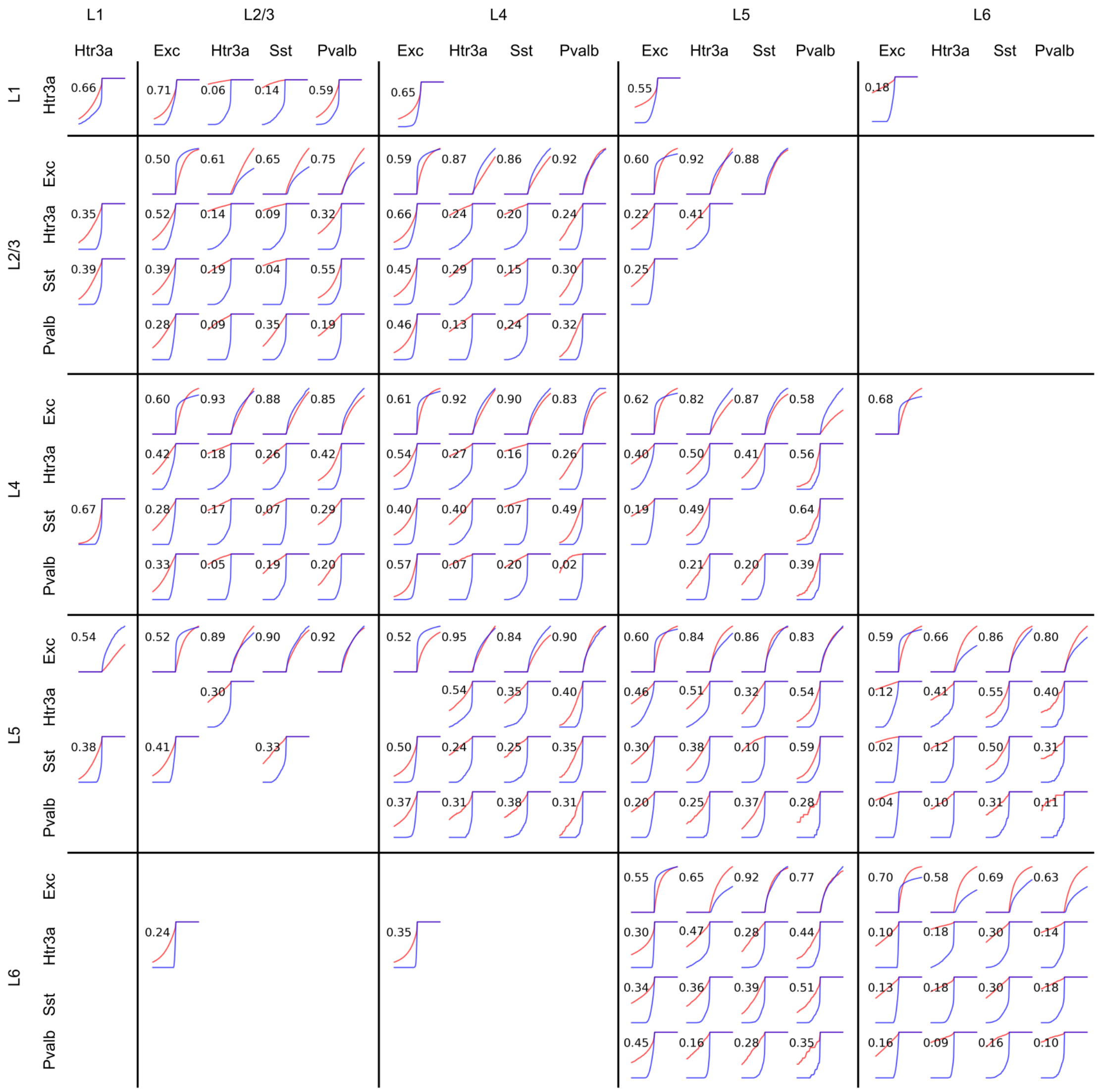


Figure S3: **Weight distributions for specific connection types.** Rows are labeled by presynaptic neuron types and columns by postsynaptic neuron types. Cumulative weight distributions are shown before (blue) and after training (red). Numbers indicate for each pair of distributions their similarity score according to Billeh et al. (2020).

Siegle, Joshua H., Xiaoxuan Jia, Séverine Durand, Sam Gale, Corbett Bennett, Nile Graddis, Greggory Heller, Tamina K. Ramirez, Hannah Choi, Jennifer A. Luviano, Peter A. Groblewski, Ruweida Ahmed, Anton Arkhipov, Amy Bernard, Yazan N. Billeh, Dillan Brown, Michael A. Buice, Nicolas Cain, Shiella Caldejon, Linzy Casal, Andrew Cho, Maggie Chvilicek, Timothy C. Cox, Kael Dai, Daniel J. Denman, Saskia E. J. de Vries, Roald Dietzman, Luke Esposito, Colin Farrell, David Feng, John Galbraith, Marina Garrett, Emily C. Gelfand, Nicole Hancock, Julie A. Harris, Robert Howard, Brian Hu, Ross Hytnen, Ramakrishnan Iyer, Erika Jessett, Katelyn Johnson, India Kato, Justin Kiggins, Sophie Lambert, Jerome Lecoq, Peter Ledochowitsch, Jung Hoon Lee, Arielle Leon, Yang Li, Elizabeth Liang, Fuhui Long, Kyla Mace, Jose Melchior, Daniel Millman, Tyler Mollenkopf, Chelsea Nayan, Lydia Ng, Kiet Ngo, Thuyahn Nguyen, Philip R. Nicovich, Kat North, Gabriel Koch Ocker, Doug Ollerenshaw, Michael Oliver, Marius Pachitariu, Jed Perkins, Melissa Reding, David Reid, Miranda Robertson, Kara Ronellenfitch, Sam Seid, Cliff Slaughterbeck, Michelle Stoecklin, David Sullivan, Ben Sutton, Jackie Swapp, Carol Thompson, Kristen Turner, Wayne Wakeman, Jennifer D. Whitesell, Derric Williams, Ali Williford, Rob Young, Hongkui Zeng, Sarah Naylor, John W. Phillips, R. Clay Reid, Stefan Mihalas, Shawn R. Olsen, and Christof Koch (Jan. 2021). “Survey of spiking in the mouse visual system reveals functional hierarchy”. In: *Nature*. ISSN: 1476-4687. DOI: 10.1038/s41586-020-03171-x. URL: https://doi.org/10.1038/s41586-020-03171-x.
